# The Cumulative Distribution Function Based Method for Random Drift Model

**DOI:** 10.1101/2025.09.29.679385

**Authors:** Chenghua Duan, Chun Liu, Xingye Yue

## Abstract

We propose a numerical method to uniformly handle the random genetic drift model for pure drift with or without natural selection and mutation. For pure drift and natural selection case, the Dirac δ singularity will develop at two boundary ends and the mass lumped at the two ends stands for the fixation probability. For the one-way mutation case, known as Muller’s ratchet, the accumulation of deleterious mutations leads to the loss of the fittest gene, the Dirac δ singularity will spike only at one boundary end, which stands for the fixation of the deleterious gene and loss of the fittest one. For two-way mutation case, the singularity with negative power law may emerge near boundary points. We first rewrite the original model on the probability density function (PDF) to one with respect to the cumulative distribution function (CDF). Dirac δ singularity of the PDF becomes the discontinuity of the CDF. Then we establish a revised finite difference method(rFDM), which keeps the total probability, is positivity-preserving and unconditionally stable. For pure drift, the scheme also keeps the conservation of expectation. It can catch the discontinuous jump of the CDF, then predicts accurately the fixation probability for pure drift with or without natural selection and one-way mutation. For two-way mutation case, it can catch the power law of the singularity. The numerical results show the effectiveness of the scheme. We also compare rFDM with a standard finite difference method(sFDM). We find that, for pure drift problem, sFDM fails to keeps the conservation of expectation and can not predict the fixation probability, because there is artificial mutation introduced near two boundary ends.

## 1 Introduction

In the population genetics, the random genetic drift model describes that the number of gene variants (alleles) fluctuates randomly over time due to random sampling. The fraction of an allele in the population can be used to measure the intensity of the random genetic drift. In other words, the value of the fraction equals to zero or one, which means the allele disappears or is completely chosen in the system. Population genetics models, aiming at modeling genetic variability, had a natural start with discrete stochastic models at the individual level. The earlier mathematical model of random genetic drift, known as the Wright-Fisher model, was constructed by Ronald Fisher [10] and Sewall Wright [23–25]. The Wright-Fisher model is regarded as a discrete-time Markov chain under the assumption that the generations do not overlap and that each copy of the gene of the new generation is selected independently and randomly from the whole gene pool of the previous generation. Then Moran [17,18] and Kimura [12,14–16] derived the diffusion limit of random Wright-Fisher model. Chalub et. al spreaded the large population limit of the Moran process, and obtained a continuous model that may highlight the genetic-drift (neutral evolution) or natural selection [3, 15, 20].

We consider a population of constant size *N*_*e*_ with a pair of two types A and B. Time *t* is increased by the time step Δ*t*, and the process is repeated. *x* is the gene *A* frequency at *t* generation. Let 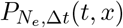 denote the probability of finding a fraction *x* of type *A* individuals at time *t* in population of fixed size *N*_*e*_ evolving in discrete time, according to the Moran process. Then, in the limit of large population and small time steps, we postulate the existence of a probability density of states, that will depend on the precise way the limits are taken [3], namely,

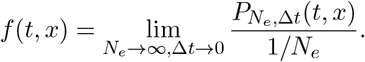

In this paper, we study the following degenerated Fokker-Planck equation on the probability density function (PDF) *f* (*t, x*):

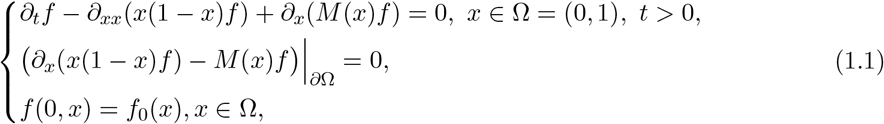

where *f*_0_(*x*) is the initial PDF and the function *M* (*x*), which is typically a polynomial in *x*, incorporates the forces of mutation and selection [15, 16, 28].

Eq.(1.1) satisfies the following conservation laws:

a. Mass conservation law:

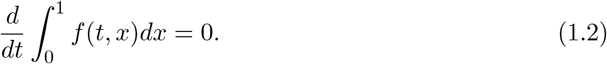
b. Moment conservation law:

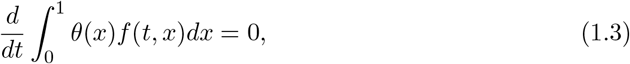

where *θ*(*x*) is the fixation probability function that satisfies

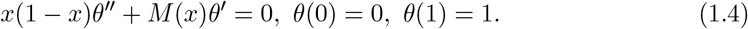

One notes that (1.3) recovers the conservation of expectation for pure drift with *θ*(*x*) = *x*. Based on different *M* (*x*), the behavior of the solution has three cases:

### Case 1. Pure drift and natural selection

For pure drift, we have *M* (*x*) = 0. When natural selection is involved, we have [2, 4],

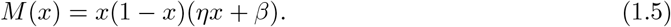

The important theoretically relevant results were shown in [2–4]. Let ℬ ℳ^+^([0, 1]) denote the space of (positive) Radon measures on [0, 1].

#### Definition 1.1.

*A weak solution to* (1.1) *is a function in L*^∞^([0, ∞), ℬ ℳ^+^([0, 1])) *satisfying*

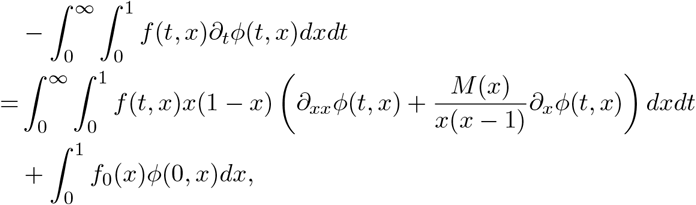

*for any* 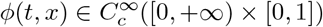.

The following theorem can be found in [4].

#### Theorem 1.2.

*If f*_0_ ∈ ℬ ℳ^+^([0, 1]) *and M* (*x*) = 0 *or given by* (1.5), *then there exists a unique weak solution f* ∈ *L*^∞^([0, ∞); ℬ ℳ^+^([0, 1])) ∩ *C*^∞^(ℝ^+^, *C*^∞^(0, 1)) *to* *equation* (1.1), *in a sense of Definition* (1.1), *such that the conservation laws* (1.2)*-*(1.3) *are valid. The solution has a form as*

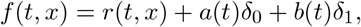

*where r*(*t, x*) ∈ *C*^∞^(ℝ^+^; *C*^∞^([0, 1])) *is a classical solution to* (1.1) *without any boundary condition*, δ_*s*_ *denotes the singular point measure supported at s, functions a*(*t*) *and b*(*t*) ∈ *C*([0, ∞)) ∩ *C*^∞^(ℝ^+^). *Furthermore, as t* → ∞, *r*(*t, x*) → 0 *uniformly. And a*(*t*) *and b*(*t*) *are monotonically increasing functions such that*

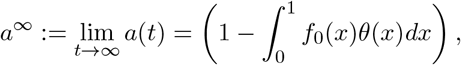

*and*

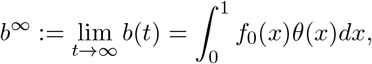

*where θ*(*x*) *is given by* (1.4). *Finally, the convergence rate is exponential*.

The appearance of the point measure δ_0_ (δ_1_) stands for that the fixation at gene *B* (*A*) happens with a probability *a*(*t*) (*b*(*t*)). The spatial temporal dynamics of the Kimura equation are well understood in the purely diffusive case and in only a relatively small number of population [5,6,13].

### Case 2. One-way mutation: Muller’s ratchet

Considering the one-way mutation from gene *B* to gene *A*,

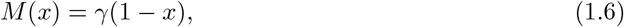

where the constant *γ >* 0 stands for the mutation rate [8, 11, 16, 21].

A well-known model is Muller’s ratchet, i.e, the fittest gene *B* of individuals is eventually lost from the population and deleterious mutations (*B* → *A*) slowly but irreversibly accumulate through rare stochastic fluctuations [9, 19]. In a finite asexual population, offspring inherit all the deleterious mutations their parents possess. Since these offspring also occasionally acquire new deleterious mutations, populations will tend to accumulate deleterious mutations over time. This effect is known as Muller’s ratchet.

There exists a unique steady solution *f*_∞_,

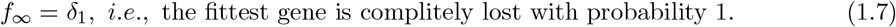

### Case 3. Two-way mutation

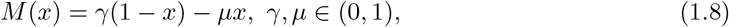

where the constant *µ >* 0 stands for the mutation rate of gene *A* to gene *B* and *γ >* 0 is the rate in opposite direction. In the long term one might expect to exist an equilibrium state due to the two direction mutation. Actually, there exists a unique steady state solution *f*_∞_(*x*) to (1.1) and (1.8),

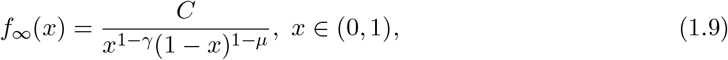

where *C* is the constant such that 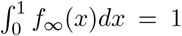. Only singularity of negative power law develops at the ends, rather than Dirac δ appears for the cases of pure drift, natural selection and one-way mutation.

For the numerical simulation, the crucial features to solve (1.1) are that the numerical solution can keep the total probability conservation law and accurately capture the concentration phenomena at the discrete level. The total probability fails to keep the total probability by some classical numerical schemes [1, 16, 22]. A complete solution, whose total probability is unity, obtained by finite volume method (FVM) schemes in [28]. Xu et al. [26] compared a serial of finite volume and finite element schemes for the pure diffusion equation. Their critical comparison of the long-time asymptotic performance urges carefulness in choosing a numerical method for this type of problem, otherwise the main properties of the model might be not kept. In recent years, a variational particle method was proposed based on an energetic variational approach, by which a complete numerical solution can be obtained and the positivity of the solution can be kept [7]. However, some artificial criteria must be introduced to handle the concentrations near the boundary ends, even though it is designed based on the biological background. Recently, an optimal mass transport method based on pseudo-inverse of CDF is used to solve the model [2]. In this method, the feasible solution is strictly monotonous. However, for the cases of pure drift, natural selection and one-way mutation, the Dirac-δ concentrations must be developed, then the corresponding CDF must be discontinuous at the concentration points. This leads to a fact that its pseudo-inverse can not be strictly monotonous. However, numerical results were presented for the cases of pure drift and selection. So some manual intervention must be introduced in the numerical implements. The pure drift problem with multi-alleles, corresponding to a multidimensional PDE, is investigated in [27] by finite-difference methods, where the authors proposed a numerical scheme with absolute stability and several biologically/physically relevant quantities conserved, such as positivity, total probability, and expectation. So far, although quite a few numerical methods have been established for the pure drift and selection cases, efficient numerical works on the mutation case are not reported yet.

In this paper, we rewrite the system (1.1) to the following new one on the cumulative distribution function (CDF) 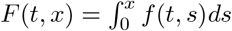 as

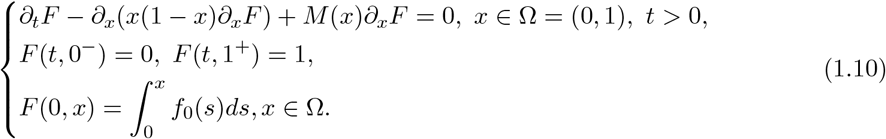

Taking the CDF into account, the Dirac δ singularity at the boundary points of original PDF *f* will change to the discontinuity of the CDF *F* at the boundary points, and the fixation probability (lumped mass) will change to the height of the discontinuous jump. The singularity of negative power law will change to a bounded positive power law.

We will propose a finite difference scheme for (1.10) with a key revision near the boundary, which can handle the pure drift with or without selection and mutation, is unconditional stable, and keeps the total probability and positivity. It also keeps the conservation of expectation for pure drift. The numerical results show that the scheme can catch the height of discontinuity at the ends and predict accurately the fixation probability for the cases of pure drift, natural selection and one-way mutation. It also predict accurately the negative power of the power law for two-way mutation case.

The rest of this paper is organized as follows. In Section 2, we construct the numerical scheme for (1.10). Some numerical analysis is presented in Section 3. In Section 4, several numerical examples are presented to validate the theoretical results and to demonstrate the ability to trace the long-time dynamics of random genetic drift. Some discussions will be presented in Section 5 about the relations between the revised scheme and the standard finite difference method. We find that the standard finite difference method does not work since an artificial mutation term is introduced.

## 2 Numerical Scheme

In this section, we introduce the revised finite difference method (rFDM) for (1.10) with the central difference method for diffusion term and the upwind scheme for convection term. Let *h* = 1*/K* be the spatial step size, and *x*_*i*_ = *ih, i* = 0, · · ·, *K*, be the spatial grid points. Let *τ* be the temporal step size, and *t*_*n*_ = *nτ, n* = 1, 2, · · ·, *N*, be the temporal grid points.

Define the first order difference as

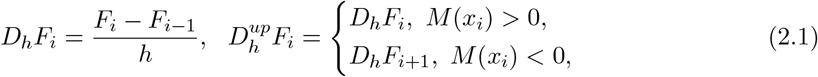

and denote the diffusion coefficient by *a*(*x*) = *x*(1 − *x*). The upwind numerical scheme, referred as **rFDM**, reads as: Given *F* ^*n*−1^, *F* ^*n*^ solves the following linear algebra system,

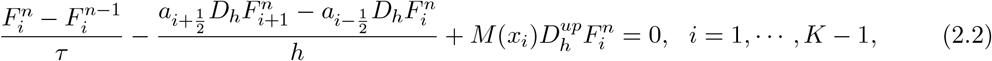

with 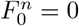 and 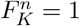, for *n* = 1, · · ·, *N*, and the key revision on the diffusion coefficient

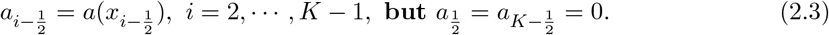

Finally, the solution of the original equation (1.1) can be recovered from *F* by, for *n* = 1, · · ·, *N*,

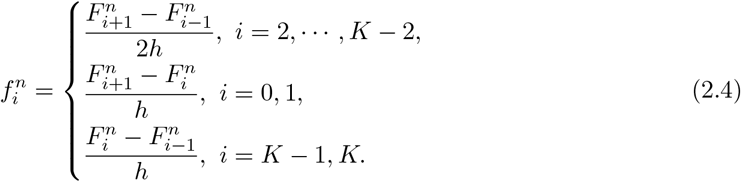

In the above formula, central difference does not applied at near boundary points *i* = 1 or *K* − 1, thanks to the fact that the discontinuous jump may occur at the boundary points.

### Remark 2.1

*For problem* (1.10) *in continuous sense, the diffusion coefficient a*(0) = *a*(1) = 0 *degenerates at the boundary points. This means that the information at the boundary points can never be transferred into the domain* Ω = (0, 1) *by diffusion. In our revised treatment* (2.3) *for numerical scheme, we set* 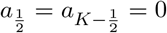, *where a term of O*(*h*) *order is omitted, since the exact value* 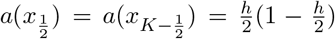. *With this revised treatment, the boundary value* 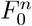 *and* 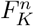*is not involved in the discrete system* (2.2) *if M* ≡ 0, *i*.*e*., *the boundary value can never be transferred into the inner points by discrete diffusion. In Section 5, we will discuss the standard scheme without this revision*.

## 3 Analysis Results

In this section, we will focus on the numerical analysis for rFDM (2.2), including the unconditional stability, positivity preserving, and conservation law of the total probability and expectation.

### Theorem 3.1

*The upwind scheme rFDM* (2.2) *is unconditionally stable and positivity-preserving. Proof*. Without loss of generality, assume there exists *i*^∗^ such that *M* (*i*) *>* 0, *i* = 1, · · ·, *i*^∗^ and *M* (*i*) *<* 0, *i* = *i*^∗^ + 1, · · ·, *K* − 1. (2.2) can be written as:

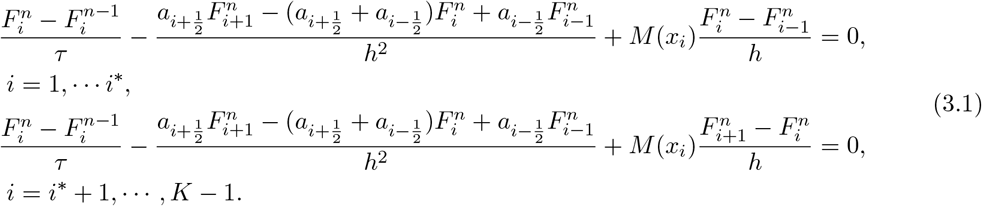

with 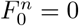, 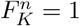 at time *t*^*n*^, *n* = 1, · · ·, *N*.

Let **B** = (*b*_*ij*_) be the matrix of the linear system. Then **B** is a tri-diagonal matrix. For *i* = 1, · · ·, *i*^∗^,

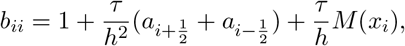

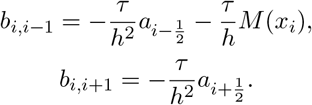

For *i* = *i*^∗^ + 1, · · ·, *K* − 1,

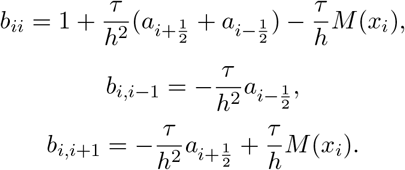

For *i* = 0, *b*_*ii*_ = 1, *b*_*i*,*i*+1_ = 0. For *i* = *K, b*_*ii*_ = 1, *b*_*i*,*i*−1_ = 0.

Note that

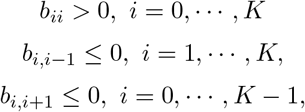

And

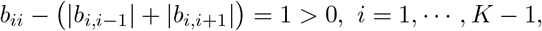

Then **B** is a M-matrix, i.e., any entry of the inverse matrix **B**^−1^ is positive. So rFDM (2.2) is unconditionally stable and positivity-preserving.

### Theorem 3.2

*The numerical scheme rFDM* (2.2) *keeps the conservation of total probability. For pure drift case, the conservation of expectation also holds*.

*Proof*. Define 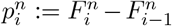 be the probability for the fraction of gene *A* belongs to the interval *I*_*i*_ = [*x*_*i*−1_, *x*_*i*_]. Then the total probability is

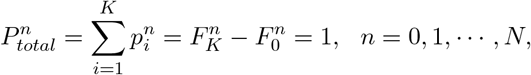

i.e., the discrete system (2.2) keeps the mass conservation naturally.

By the integral by parts, the expectation ℰ (*t*) satisfies that

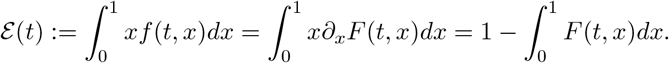

So we define a discrete expectation as

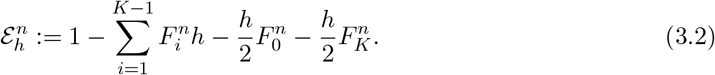

For pure drift case, *M* (*x*) ≡ 0, then we have, by (2.2) and (2.3), that

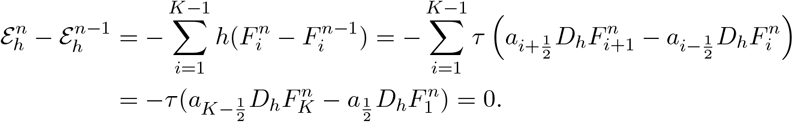

So the discrete system (2.2) keeps the conservation of the discrete expectation for pure drift case. Please note that, without the revised treatment (2.3), the conservation of the discrete expectation is invalid.

## 4 Numerical results

In this section, we will show the effectiveness of this algorithm by different numerical examples. In Example 1, we verify the local convergence. In Section 4.1, the pure drift *M* = 0 and natural selection *M* (*x*) = *x*(1 − *x*)(*ηx* + *β*) case are studied in Example 2 and Example 3, respectively.In Section 4.2, we consider the mutation case *M* (*x*) = *γ*(1 − *x*) − *µx*, including one-way mutation model, such as the Muller’s ratchet model in Example 4 and two-way mutation model in Example 5.

### Example 1. Local convergence

Although the singularity may develop at boundary points, the solution in interior area is sufficiently smooth. To verify the correctness of the numerical scheme, we check the local convergence.

Let Ω^*′*^ ⊂ Ω be the interior area and

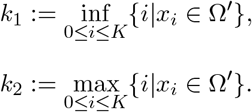

Define the error *e* of *F* (*t, x*) in ℒ^2^(Ω^*′*^) and ℒ^∞^(Ω^*′*^) mode as

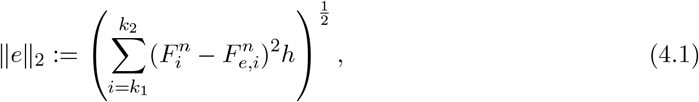

And

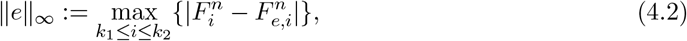

where 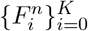is the numerical solution of CDF model (1.10), and 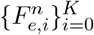 is the corresponding exact solution at time *t*^*n*^, *n* = 1, · · ·, *N*.

In this example, we take the initial probability density as

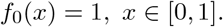

Table 1 shows the error and local convergence order of *F* (*t, x*) for *M* (*x*) = *x*(1 − *x*)(−4*x* + 2) and *M* (*x*) = 0.2(1 − *x*) + 0.4*x* in ℒ^2^([0.3, 0.7]) and ℒ^∞^([0.3, 0.7]) mode with different space and time grid sizes at time *t* = 0.1. The reference “exact” solution is obtained numerically on a fine mesh with *h* = 1*/*100000 and *τ* = 1*/*100000. The results show that the local convergence of the numerical scheme is 2nd order in space and 1st order in time for different *M* (*x*) in the inner region [0.3, 0.7].

**Table 1:**
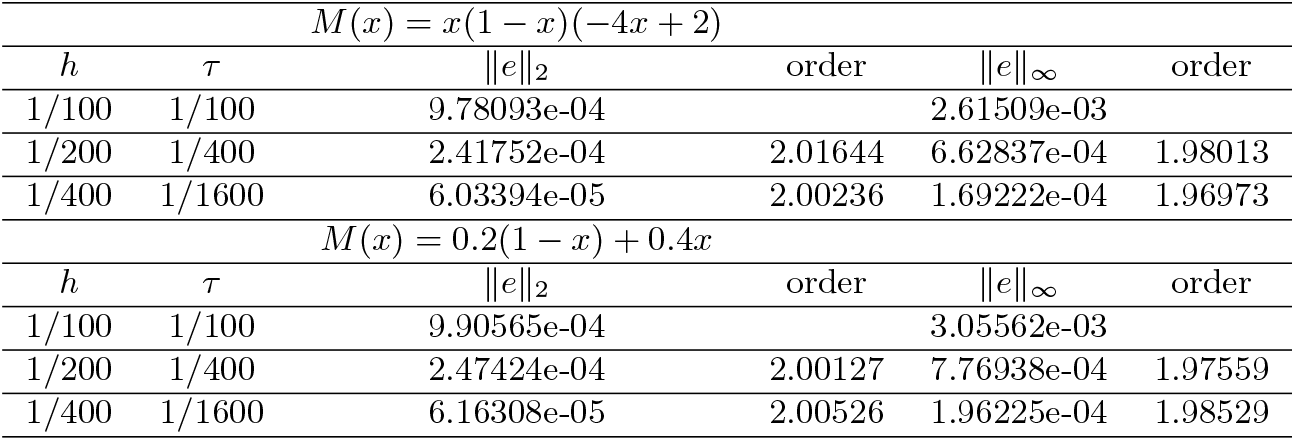
Error of *F* in Ω^*′*^ = [0.3, 0.7] at *t* = 0.1 in **Example 1**.

### 4.1 Pure drift and natural selection

In this section, we study pure drift and natural selection case which keeps the conservation law (1.2) and (1.3).

Firstly, we define the error of fixation probability at the left and right boundary points at large enough time (near the steady state) *T* = *τ N* as

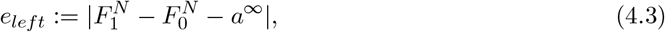

And

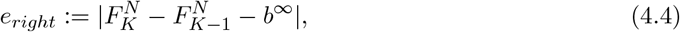

where *a*^∞^ and *b*^∞^ are the exact fixation probability at boundary points given in Theorem 1.2.

#### Example 2. Pure drift

In this example, we consider pure drift case, i.e., *M* (*x*) = 0, with a Gaussian distribution initial function as

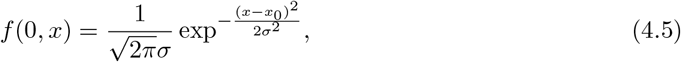

with *σ* = 0.01 and *x*_0_ = 0.7.

In Fig.1, the evolution of CDF *F* (*t, x*) are shown with partition *h* = 1*/*1000, *τ* = 1*/*1000. The discontinuity seems to develop at the boundary as time evolves. That means Dirac δ singularities develop at the boundary points for the original PDF *f*. To verify that the scheme can catch the height of the discontinuous jump, i.e., the fixation probability, we present the results in Table 2 on different spacial grid sizes (*h* = 1*/*100, 1*/*200, 1*/*400, 1*/*800) with a very fine time step *τ* = 1*/*10000 at sufficiently large time *T* = 36. It can be found that the discontinuity occurs at boundary points and the height of the jump on the two ends approach to the fixation probability given in Theorem 1.2 with a rate of 1st order. In Fig.2, the discrete expectation in (3) keeps conservation as time evolves with *h* = 1*/*1000, *τ* = 1*/*1000.

**Figure 1.**
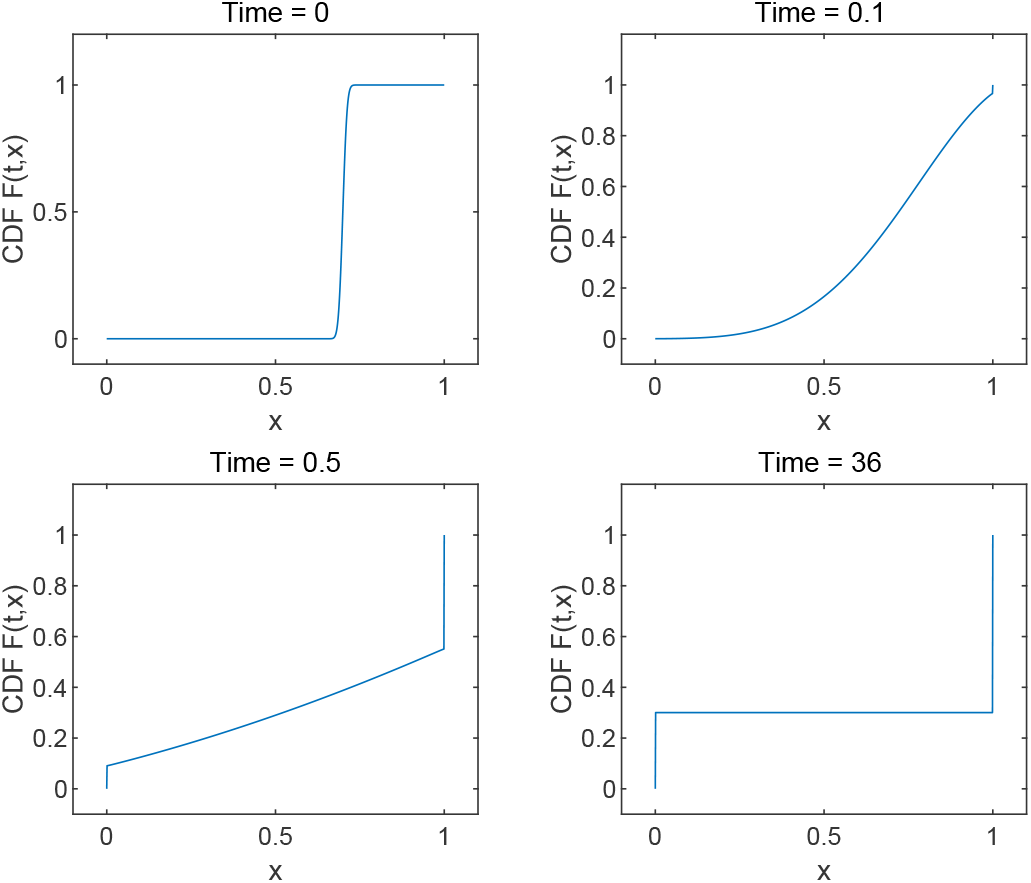
Evolution of *F* (*t, x*) in **Example 2** with *h* = 1*/*1000, *τ* = 1*/*1000.

**Figure 2.**
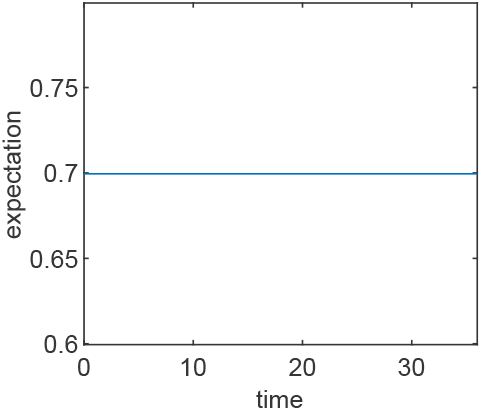
Evolution of expectation in **Example 2** with *h* = 1*/*1000, *τ* = 1*/*1000.

**Table 2:** Discontinuity at the boundary points at *T* = 36 with *τ* = 1*/*10000 in **Example 2**.

| $h$ | $F_1^N - F_0^N$ | $e_{left}$ | Order | $F_K^N - F_{K-1}^N$ | $e_{right}$ | Rate |
| --- | --- | --- | --- | --- | --- | --- |
| 1/100 | 0.303030 | 3.03030e-03 |  | 0.696970 | 3.03030e-03 |  |
| 1/200 | 0.301508 | 1.50754e-03 | 1.00727 | 0.698492 | 1.50753e-03 | 1.00727 |
| 1/400 | 0.300752 | 7.51880e-04 | 1.00362 | 0.699248 | 7.51880e-04 | 1.00362 |
| 1/800 | 0.300375 | 3.75469e-04 | 1.00181 | 0.699624 | 3.75469e-04 | 1.00180 |

<sup>1</sup> $a^\infty = 1 - x_0 = 0.3$ , $b^\infty = x_0 = 0.7$ in (4.3)-(4.4).

#### Example 3. Natural selection

In this example, we consider the natural selection case:

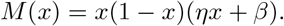

Let

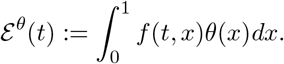

The conserved quantity (1.3) is ℰ ^*θ*^ (*t*) = ℰ ^*θ*^ (0) with

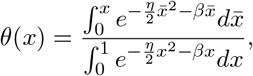

being the solution of (1.4).

By the integration by parts, we have

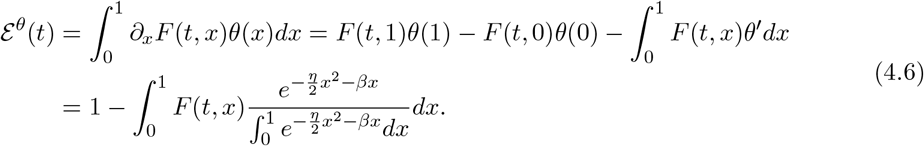

Then a discrete conserved quantity can be defined as

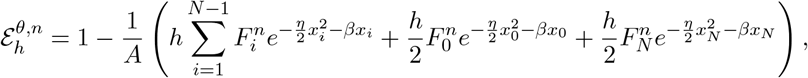

where *A* is an approximation of 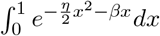,

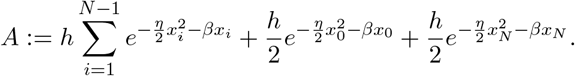

In this example, *η* = −4 and *β* = 2 and the initial state is given in (4.5) with *x*_0_ = 0.7 and *σ* = 0.01. Fig.3 shows that the discontinuous points emerge at the boundary points. As time evolves, the discrete expectation 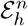does not keep the conservation and tends to a certain value (≈ 0.671595) in Fig.4, but 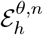keeps the conservation and its value is about 0.671529 in Fig.5. Table 3 shows the ability to catch the jump of the discontinuity at boundary points under different spacial grid sizes (*h* = 1*/*100, 1*/*200, 1*/*400, 1*/*800), and a very fine time step *τ* = 1*/*10000 at *T* = 15 when the steady state is approaching. The fixation probability is predicted in a 1-order accuracy at left and right boundary point.

**Figure 3.**
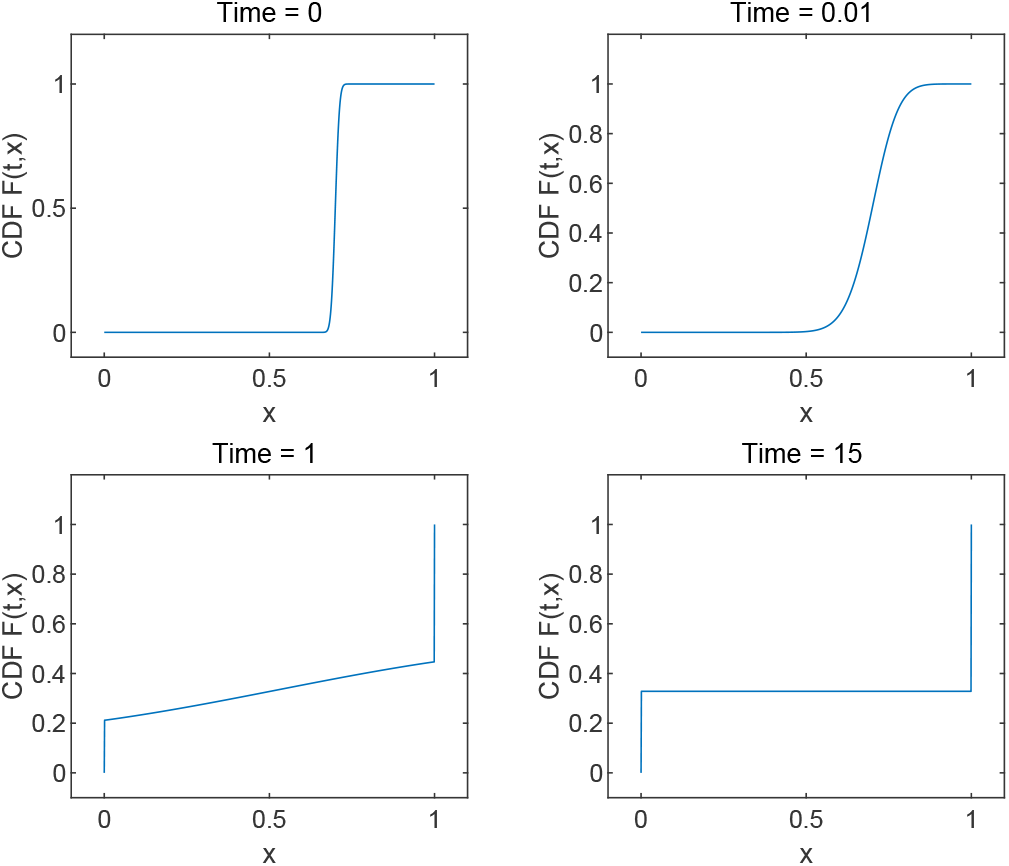
Evolution of the numerical solution of CDF in **Example 3** with *h* = 1*/*1000, *τ* = 1*/*1000.

**Figure 4.**
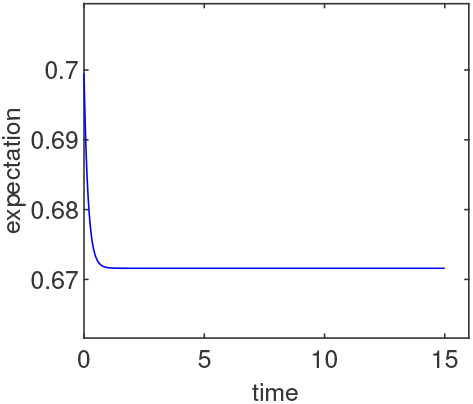
Evolution of expectation.

**Figure 5.**
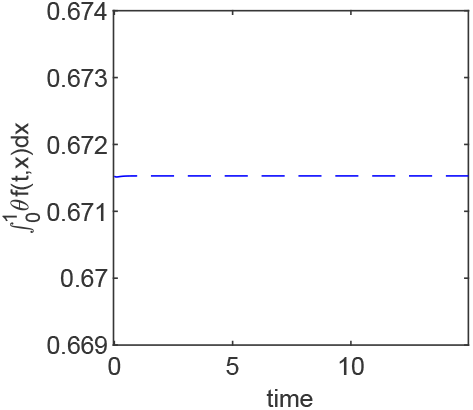
Evolution of 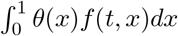.

**Table 3:** Discontinuity at the boundary points at *T* = 15 with *τ* = 1*/*10000 in **Example 3**.

| $h$ | $F_1^N - F_0^N$ | $e_{left}$ | Rate | $F_K^N - F_{K-1}^N$ | $e_{right}$ | Rate |
| --- | --- | --- | --- | --- | --- | --- |
| 1/100 | 0.330230 | 2.16298e-03 |  | 0.669770 | 2.16298e-03 |  |
| 1/200 | 0.329103 | 1.03683e-03 | 1.06085 | 0.670896 | 1.03683e-03 | 1.06085 |
| 1/400 | 0.328556 | 4.89322e-04 | 1.08332 | 0.671444 | 4.89322e-04 | 1.08332 |
| 1/800 | 0.328287 | 2.20144e-04 | 1.15234 | 0.671713 | 2.20144e-04 | 1.15234 |

<sup>1</sup> $e_{left}$ and $e_{right}$ are defined in (4.3) and (4.4), with $b^\infty = \mathcal{E}_{h^*}^{\theta,0}$ and $a^\infty = 1 - b^\infty$ , where the conserved quantity $\mathcal{E}_{h^*}^{\theta,0} = 0.671933$ under a very fine space step $h = 1/10000$ .

### 4.2 Mutation case

In this section, we discuss the mutation case *M* (*x*) = *γ*(1 − *x*) − *µx, µ, γ* ≥ 0, including one-way mutation and two-way mutation.

#### Example 4. One-way mutation: Muller’s ratchet

One-way mutation *M* (*x*) = *γ*(1 − *x*) with *γ* = 0.2 is considered in this example. The initial state is taken as

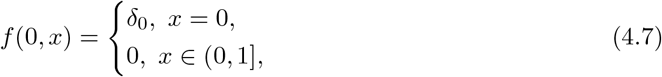

i.e., there is a point measure with the whole probability 1 at *x* = 0 at the initial time. That means only the fittest gene *B* exists in the system. Fig.6 shows the evolution of CDF *F* (*t, x*). The discontinuity develops at *x* = 1 and the height of the discontinuous jump rises up to 1 eventually. Table 4 shows the numerical results with different spatial step sizes (*h* = 1*/*100, 1*/*200, 1*/*400, 1*/*800) and a very fine time step *τ* = 1*/*10000 at *T* = 50, when the steady state is approaching. It can be found that the discontinuity only emerges at *x* = 1 with height of 1 and no discontinuity happens at *x* = 0. This fact accords with the Muller’s ratchet theory: all fittest gene *B* will mutate irreversibly to the deleterious gene *A*.

**Figure 6.**
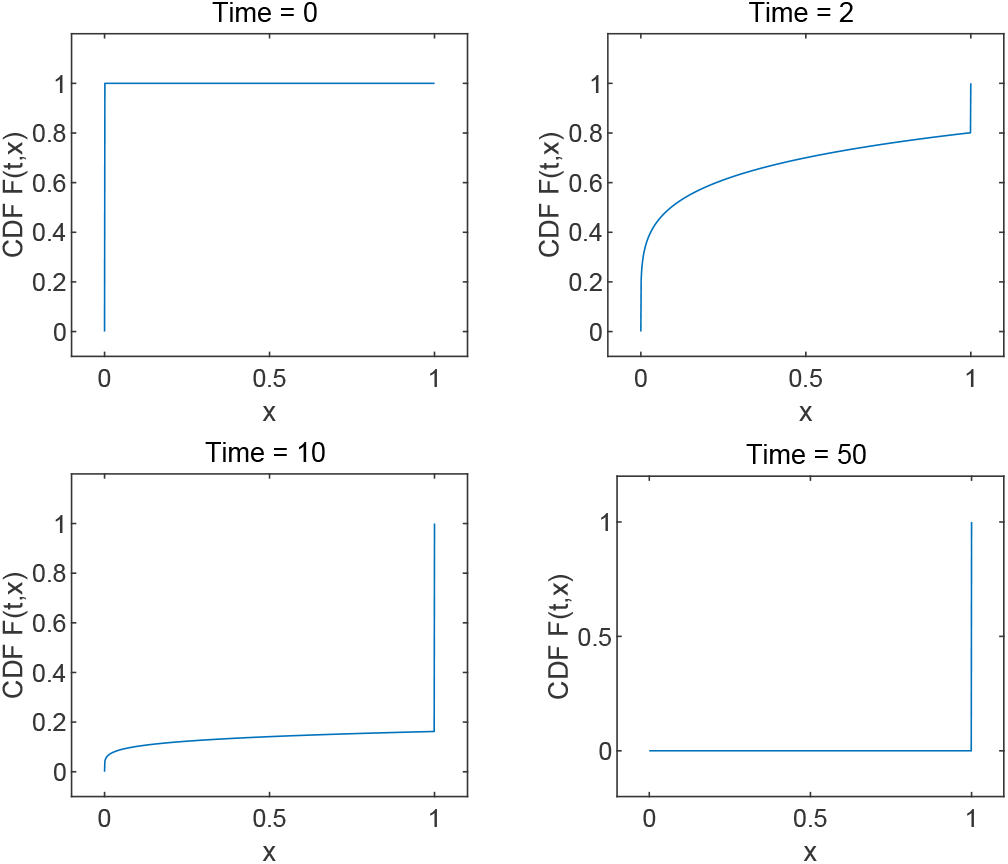
Evolution of *F* (*t, x*) in **Example 4** for *γ* = 0.2, *µ* = 0 with *h* = 1*/*1000, *τ* = 1*/*1000.

**Table 4:** Discontinuity at the right boundary point at *T* = 50 in **Example 4**.

| $h$ | $F_1^N - F_0^N$ | $F_K^N - F_{K-1}^N$ |
| --- | --- | --- |
| 1/100 | 2.22716e-05 | 0.999946 |
| 1/200 | 1.93788e-05 | 0.999946 |
| 1/400 | 1.68642e-05 | 0.999946 |
| 1/800 | 1.46777e-05 | 0.999946 |

#### Example 5. Two-way mutation

In this example, we consider a two-way mutation *M* (*X*) = *γ*(1−*x*)−*µx* with *µ* = 0.2, *γ* = 0.4. Taking into account that *F* (*t, x*) may be smooth on *x* ∈ [0, 1], the probability density *f* (*t, x*) can be recovered by (2.4) except *i* = 1, *K* − 1. Central difference is available now,

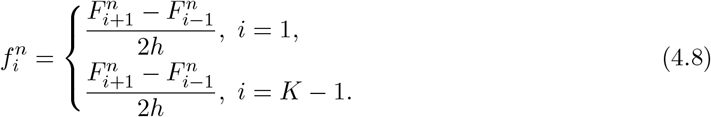

The initial function is chosen as (4.5) with *σ* = 0.01. Fig.7 shows that the CDF may be continuous as time evolves with *x*_0_ = 0.7 under *h* = 1*/*1000, *τ* = 1*/*1000. Fig.8 shows the expectation is not conserved as time evolves and towards to the same value 0.66677 with different initial *x*_0_ = 0.7, 0.2. The left figure of Fig.9 shows the relationship between ln(*F* (*t, x*)) and ln(*x*) is approximately linear at *T* = 36 with *h* = 1*/*3200 and *τ* = 1*/*10000. ln(1 − *F* (*t, x*)) and ln(1 − *x*) are also approximately linear in the right figure of Fig. 9. The results show that *F* (*t, x*) can be approached by polynomial with *x*^*ξ*^ near *x* = 0 and (1 − *x*)^*η*^ near *x* = 1, where *ξ, η* are positive constant, at *T* = 36 near the steady state.

**Figure 7.**
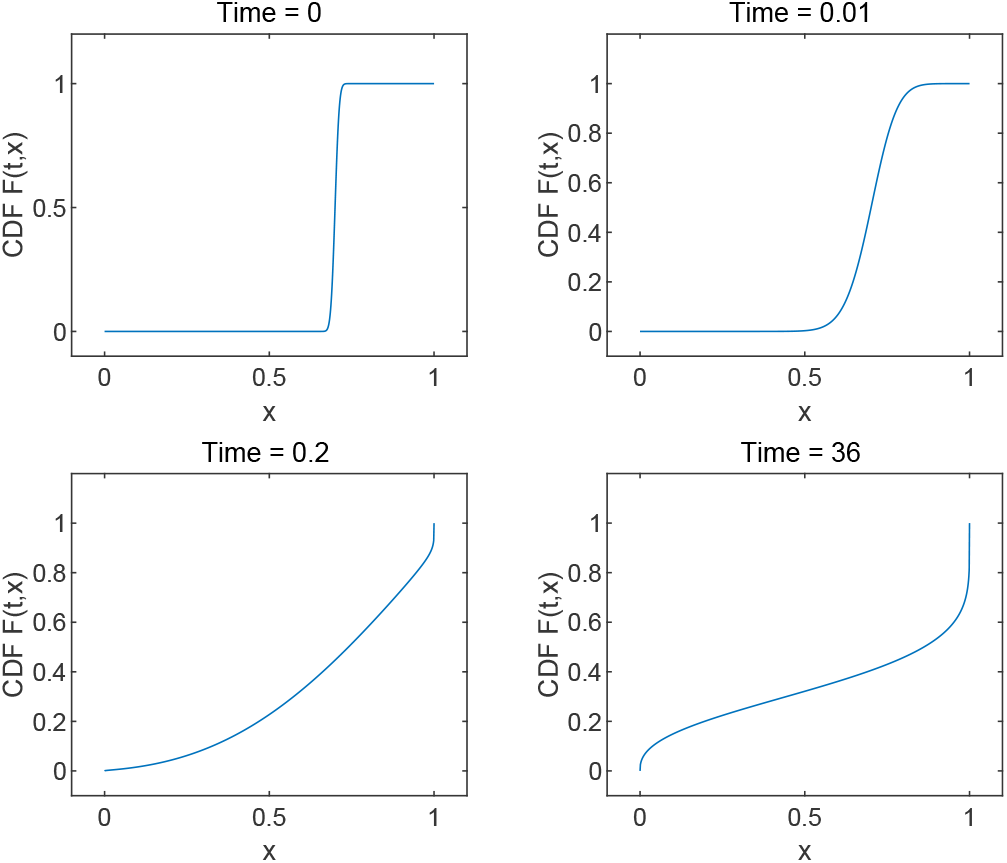
Evolution of *F* (*t, x*) in **Example 5** for *µ* = 0.2, *γ* = 0.4 with *h* = 1*/*1000, *τ* = 1*/*1000.

**Figure 8.**
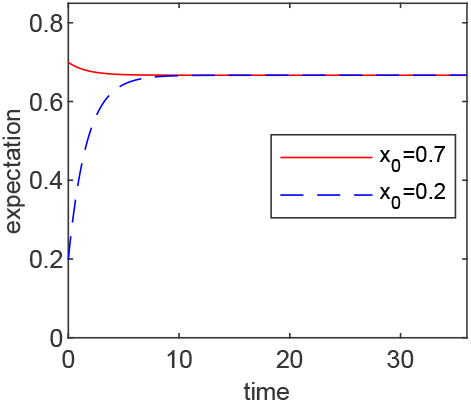
Evolution of expectation in **Example 5** with *µ* = 0.2, *γ* = 0.4 under *h* = 1*/*1000, *τ* = 1*/*1000.

**Figure 9.**
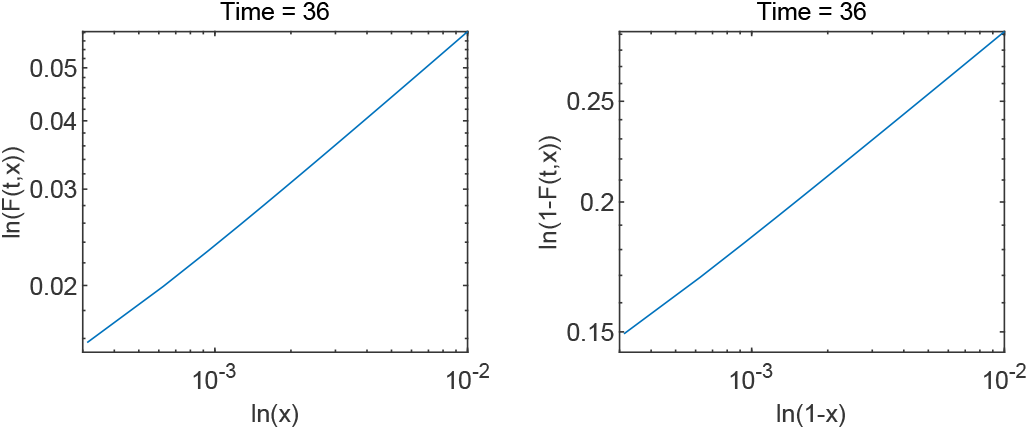
ln(*F* (*t, x*)) near boundary points in **Example 5** at *T* = 36 with *h* = 1*/*3600, *τ* = 1*/*10000.

Table 5 shows the behavior of the power law near the boundary points *x* = 0, 1 with different initial states (*x*_0_ = 0.7 and *x*_0_ = 0.2) under different spatial mesh steps and the refine time step *τ* = 1*/*10000 at time *T* = 36. The value of 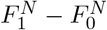 means the point measure does not emerge at the boundary point. The results also show that the numerical *F* (*t, x*) can be approached by polynomial *x*^*γ*^ with *γ* ≈ 0.4 near *x* = 0 and (1 − *x*)^*µ*^ with *µ* ≈ 0.2 near *x* = 1, respectively. That means the corresponding probability density *f* (*t, x*) is close to *x*^−0.6^ at *x* = 0 and (1 − *x*)^−0.8^ at *x* = 1. It suggests that the numerical results are consistent with theoretical results (1.9). In addition, numerical results also show the fact that the steady state has nothing with the initial states.

**Table 5:** Behavior of power law at boundary points at *T* = 36 with *τ* = 1*/*10000 in **Example 5**.

| $x_0 = 0.7$ | | | | | | |
| --- | --- | --- | --- | --- | --- | --- |
| $h$ | $F_0^N$ | $F_1^N$ | $\gamma$ | $F_{K-1}^N$ | $F_K^N$ | $\mu$ |
| 1/200 | 0.00000 | 4.78993e-02 |  | 7.37404e-01 | 1.00000 |  |
| 1/400 | 0.00000 | 3.62271e-02 | 0.402935 | 7.72496e-01 | 1.00000 | 0.206953 |
| 1/800 | 0.00000 | 2.74198e-02 | 0.401851 | 8.02473e-01 | 1.00000 | 0.203842 |
| 1/1600 | 0.00000 | 2.07643e-02 | 0.401110 | 8.28292e-01 | 1.00000 | 0.202093 |
| 1/3200 | 0.00000 | 1.57294e-02 | 0.400639 | 8.50637e-01 | 1.00000 | 0.201136 |
| $x_0 = 0.2$ | | | | | | |
| $h$ | $F_0^N$ | $F_1^N$ | $\gamma$ | $F_{K-1}^N$ | $F_K^N$ | $\mu$ |
| 1/200 | 0.00000 | 4.78993e-02 |  | 7.37404e-01 | 1.00000 |  |
| 1/400 | 0.00000 | 3.62271e-02 | 0.402935 | 7.72496e-01 | 1.00000 | 0.206953 |
| 1/800 | 0.00000 | 2.74198e-02 | 0.401851 | 8.02473e-01 | 1.00000 | 0.203842 |
| 1/1600 | 0.00000 | 2.07643e-02 | 0.401110 | 8.28292e-01 | 1.00000 | 0.202093 |
| 1/3200 | 0.00000 | 1.57294e-02 | 0.400639 | 8.50637e-01 | 1.00000 | 0.201136 |

<sup>1</sup> $\gamma$ and $\mu$ are obtained by

## 5 Some discussions about rFDM and standard FDM

In this section, we discuss what happens if the revised treatment (2.3) is not introduced. Recalling that *a*(*x*) = *x*(1 − *x*), the standard finite difference scheme, referred as **sFDM**, is

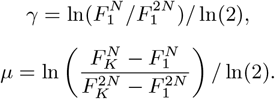

as follows. Find 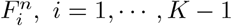 such that

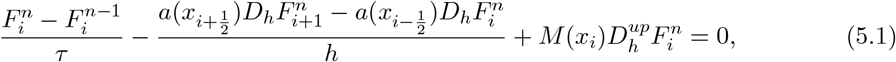

subject to 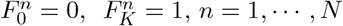.

Firstly, we compare the numerical behavior of the two schemes. Without loss of generality, we consider the pure drift case *M* (*x*) = 0 and take the initial state as (4.5) with *x*_0_ = 0.7 and *σ* = 0.01. Numerical results are presented in Figs 10 and 11.

**Figure 10.**
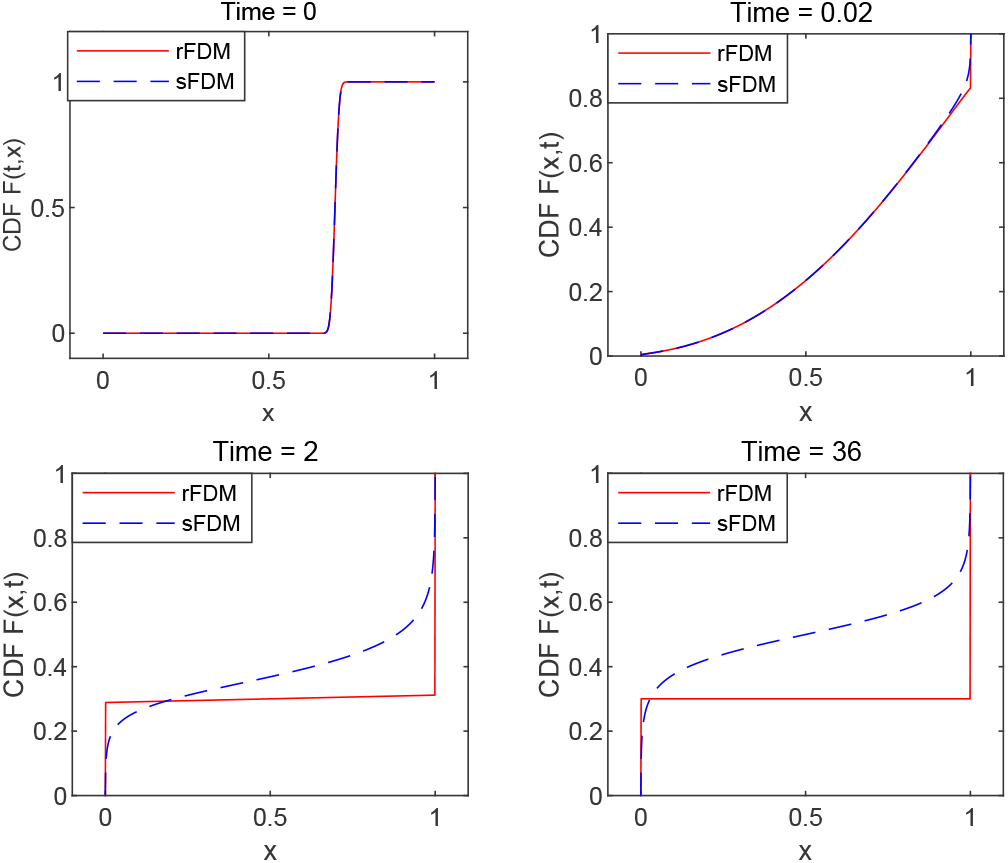
Evolution of *F* (*t, x*) by rFDM and sFDM with *h* = 1*/*1000, *τ* = 1*/*1000.

**Figure 11.**
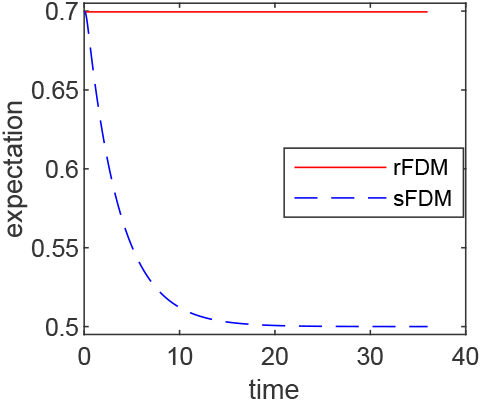
Evolution of expectation in for rFDM and sFDM with *h* = 1*/*1000, *τ* = 1*/*1000.

The evolution of CDF *F* (*t, x*) by rFDM (2.2)-(2.3) and sFDM (5.1) are shown in Fig.10. As time evolves, the discontinuity emerges at the ends *x* = 0, 1 by rFDM and the fixation phenomenon is correctly predicted. For sFDM, no evidence implies the development of the discontinuity, i.e., sFDM fails to predict the fixation phenomenon. To make it more clear, more results by sFDM are presented in Table 6 with different spatial grid size (*h* = 1*/*100, 1*/*200, 1*/*400, 1*/*800) and the fixed time step size *τ* = 1*/*10000. It is obvious that sFDM fails to catch the discontinuity that should develop at the ends.

**Table 6:** Behavior at boundary points for sFDM (5.1) at *T* = 36.

| $h$ | $F_1^N - F_0^N$ | $e_{left}$ | $F_K^N - F_{K-1}^N$ | $e_{right}$ |
| --- | --- | --- | --- | --- |
| 1/100 | 0.153003 | 0.146998 | 0.153003 | 0.546997 |
| 1/200 | 0.138051 | 0.161949 | 0.138052 | 0.561948 |
| 1/400 | 0.125864 | 0.174135 | 0.125865 | 0.574135 |
| 1/800 | 0.115703 | 0.184297 | 0.115707 | 0.584294 |

<sup>1</sup> $a^\infty = 1 - x_0 = 0.3$ , $b^\infty = x_0 = 0.7$ in (4.3)-(4.4).

The evolution of expectation in Fig. 11 shows that rFDM keeps the conservation of expectation, while sFDM fails.

The reason why sFDM does not work is that sFDM destroys the rule that the information at boundary points should not be transferred into the domain by diffusion due to the degeneration of the diffusion coefficient *a*(0) = *a*(1) = 0 at the boundary points.

Next, the result of sFDM in Fig.10 looks like the one in Fig.7 for two-way mutation case. But what we treat now is the pure drift case *M* (*x*) = 0. To make this clear, we take the difference between rFDM (2.2)-(2.3) and sFDM (5.1). The only difference takes place at two points *x*_1_ and

*x*_*K*−1_. At *x*_1_, sFDM is rFDM plus a term at the left hand side as

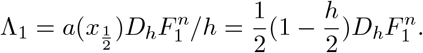

This means that at *x*_1_, a mutation from gene *B* to *A* is numerically introduced to a pure drift case, here *M* (*x*) = *γ*(1 − *x*) with a mutation ratio 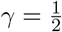. Similarly, at *x*_*K*−1_, sFDM is rFDM plus a term at the right hand side as

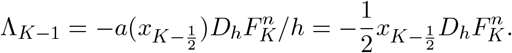

That implies that at *x*_*K* − 1_, a mutation from gene *A* to *B* is numerically introduced, here *M* (*x*) = −*µx* with a mutation rate 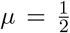. With these artificial mutations, fixation can never happen. That’s the reason why sFDM fails to predict the fixation that should happen for pure drift and its numerical results behavior like a problem with two-way mutation.

## 6 Conlusions

We re-model the random genetic drift problem on PDF to a new one on CDF. The possible Dirac δ singularity on PDF then changes to a discontinuous jump on CDF and the height of the jump is just the fixation probability. The possible singularity of negative power law changes to a bounded positive power law. A revised finite difference method is proposed to effectively handle the pure drift with or without natural selection, one-way mutation and two-way mutation in a unifying way, while standard finite difference method does not work since it introduce an artificial mutation term.

The idea working on CDF is a potential way to treat multi-alleles genetic drift problems, where multi-dimensional partial differential equations is involved. It is quite direct to change the equation on PDF to one on CDF. But it is a challenge now to settle down the boundary condition, which is corresponding to the margin distribution.

## Acknowledgments

The authors would like to thank Prof X.F. Chen for very helpful discussions on this topic.The work of the first author was supported in part by NSFC 11901109. The work of the second author is partially supported by NSF grants DMS-1950868 and DMS2118181. The work of the third author was supported by NSFC 11971342, 12071190 and 12371401.

